# Impact of Cell Size, Light Wavelengths, and Intensities on Growth, Oxygen Production, and Consumption Rates of *Chromochloris zofingiensis* and *Haematococcus lacustris*

**DOI:** 10.1101/2025.11.18.689143

**Authors:** Ugwu Chigozie, Ayumi Hashiguchi, Hideaki Nagare

## Abstract

This study examined the oxygen production, consumption, and morphological characteristics of *Chromochloris zofingiensis* (*C. zofingiensis*) and *Haematococcus lacustris* (*H. lacustris*) under autotrophic conditions using blue and red-light at different intensities (20–80 µmol m^−2^ s^−1^). Both algal species exhibited reduced growth rates as light intensity increased. At 20 µmol m^−2^ s^−1^, *C. zofingiensis* achieved a higher specific growth rate under blue-light (0.52 d^−1^) compared to red-light (0.14 d^−1^), while *H. lacustris* grew more rapidly under red-light (0.44 d^−1^) than blue-light (0.12 d^−1^). Under blue-light, *C. zofingiensis* demonstrated enhanced oxygen metabolism, with high OPR (10 µmol O_2_ gSS^−1^ min^−1^) and OCR (–7.08 µmol O_2_ gSS^−1^ min^−1^). This performance was associated with its smaller cell size (1.99–8.32 µm) and higher surface area-to-volume ratio (9.6 ± 6.4 ×10^−5^ m^2^ m^−3^). In contrast, *H. lacustris* showed lower oxygen exchange rates with OPR (5.63 µmol O_2_ gSS^−1^ min^−1^) and OCR (–4.85 µmol O_2_ gSS^−1^ min^−1^) consistent with its larger cell size (11.42–26.86 µm) and lower surface area-to-volume ratio (3.4 ± 2.1×10^−10^ m^2^ m^−3^). Both species exhibited diminished oxygen exchange under red light due to low photon energy and high pH levels. These findings underscore the influence of blue light and cell morphology on microalgae oxygen dynamics.

## 1. Introduction

Photosynthesis is the primary process by which light energy is converted into chemical energy, sustaining nearly all life on Earth. Beyond driving oxygen evolution and food chain formation, photosynthesis in aquatic microorganisms plays a particularly critical role. Microalgae are diverse group of fast-growing unicellular and multicellular microorganism. They are particularly efficient at converting light and CO_2_ into biomass [1,2] while releasing oxygen as a byproduct [3].

Marine microalgae and cyanobacteria are estimated to contribute 50–80% of the Earth’s oxygen supply and fix approximately 50 gigatons of carbon annually, surpassing terrestrial plants in both oxygen generation and carbon cycling [4,5]. This dual role underscores their ecological significance and growing potential in sustainable biotechnological applications [6,7].

In recent years, microalgal oxygen production and consumption dynamics have attracted attention in environmental engineering, wastewater treatment, and bioresource utilization [1]. Microalgae not only generate oxygen through photosynthesis but also sustain bacterial activity in algal–bacterial consortia by meeting oxygen demand during organic matter degradation [8]. Physiological indicators such as the oxygen production rate (OPR), oxygen consumption rate (OCR), and their ratio expressed as the photosynthetic quotient (PQ) are essential parameters for evaluating microalgal performance [9,10]. Together, these metrics define the microalgal net photosynthetic rate (MNPR), reflecting the balance between photosynthetic oxygen evolution and mitochondrial respiration [9]. Furthermore, oxygen plays a central role in microalgal carotenoid biosynthesis. Under high light stress, reactive oxygen species (ROS) such as singlet oxygen (¹O_2_), superoxide (O_2_ ^−^), and hydroxyl radicals (•OH) are generated, triggering antioxidative responses and enhancing astaxanthin accumulation[11]. For instance, ROS-inducing treatments have been shown to significantly increase carotenoid yields in *Dunaliella bardawil* [12]. Similarly, under heterotrophic conditions, chemically generated ROS including radical treatments induced carotenogenesis and astaxanthin to 12.58 mg L in microalgae [13]. This highlights the close interplay between light intensity, oxidative stress, oxygen metabolism, and secondary metabolite biosynthesis. Several studies have examined the role of oxygen in the biosynthesis of high value substance in microalgae from the standpoint of carotenoid accumulations. Under active photosynthesis, oxygen production exceeds its consumption. Among commercially important microalgae, *H. lacustris* and *C. zofingiensis* are notable for their ability to produce astaxanthin, a high-value carotenoid. *H. lacustris* is widely recognized as the leading natural source of astaxanthin[14], accumulating up to 1.9–7 % of its dry weight during the red cyst stage under high light stress [15]. However, its complex life cycle, low biomass yield, rigid cell walls, [16,17] and demanding light requirements (100–1500 μmol m^−2^ s^−1^) limit its large-scale application [15,18,19]. In contrast, *C. zofingiensis* is a unicellular freshwater chlorophyte that exhibits rapid growth [20], tolerance to diverse environmental conditions, flexible trophic modes (autotrophic, mixotrophic, heterotrophic), and high astaxanthin productivity (111 mg L^−1^ day^−1^) [21]. Its relatively small and fragile cells (2–17 µm) also facilitate extraction of target metabolites[22,23], making it an attractive alternative to *H. lacustris*.

Although *C. zofingiensis* and *H. lacustris* have certain advantages, few studies have directly examined how variations in cell size influence oxygen production and consumption in microalgae. Most research has focused on growth dynamics, pigment accumulation, and stress responses [24–27], leaving the relationship between morphological traits (e.g., cell size) and metabolic oxygen fluxes largely unexplored. Understanding this relationship is crucial not only for selecting species with superior oxygen productivity but also for optimizing cultivation systems for cost-effective biomass and carotenoid production.

The present study investigates the impact of cell size, different light wavelengths, and intensities on growth, OPR, OCR, MNPR and PQ in *C. zofingiensis* and *H. lacustris* under autotrophic conditions. These parameters were tracked under different light wavelengths and intensities. This work provides new insights into *C. zofingiensis* and *H. lacustris* responses to light environments and identifies physiological factors critical for optimizing microalgal cultivation.

## 2. Materials and methods

### 2.1 Algae cell

*C. zofingiensis* (synonym *Muriella zofingiensis*, culture collection No. 2175 of the National Institute for Environmental Studies, Tsukuba, Japan) and *H. lacustris* (culture collection No. 144 of National Institute for Environmental Studies, Tsukuba, Japan) were sub cultured autotrophically in a 500 mL flask with modified Bristol medium for two weeks. The medium was referred to as CZ-M1 [20] and also served as artificial wastewater. The medium consisted of the following (per litre) 0.75 g NaNO_3_; 0.175 g KH_2_PO_4_; 0.075 g K_2_HPO_4_; 0.075 g MgSO_4_·7H_2_O; 0.025 g CaCl_2_·2H_2_O; 0.025 g NaCl; 5 mg FeCl_3_; 0.287 mg ZnSO_4_·7H_2_O; 0.169 mg MnSO_4_·H_2_O; 0.061 mg H_3_BO_3_; 0.0025 mg CuSO_4_·5H_2_O; 0.00124 mg (NH_4_)_6_Mo_7_O_24_·7H_2_O and pH 6.8–7.0. The culture was continuously supplied with filtered air and the temperature was maintained at 25 °C, controlled by regulating the air temperature in the chamber. Additionally, it was illuminated with white light, Toshiba FL40SS, 37 watts (20 μmol m^−2^ s^−1^, 12L/12D). The algae cells from the culture were finally prepared for the microscopic observation of algae cells and algae oxygen dynamics experiments.

### 2.2 Culture experimental set-ups

The microscopic observation of algae cells was developed to determine the algae cell diameter and volume to surface area while at their exponential growth stage. While the algae oxygen dynamics was designed to determine the algae oxygen production and consumption rates under different light wavelengths and intensities. Algal illumination was achieved using light-emitting diodes (LEDs) at two specific wavelengths: 470-nm blue-light and 660-nm red-light. The primary objective of this study was to investigate the oxygen production and consumption rates of *C. zofingiensis* and *H. lacustris* under different light wavelengths and intensities with regards to their cell characteristics. The light wavelengths applied to measure the algae oxygen dynamics could provide sufficient photons to drive photosynthesis, yet not so intense that it damages photosystems (e.g., PSII) or cause photobleaching of pigments like chlorophyll and carotenoids [7,28].

#### i. Microscopic observation of algae cells

Sequel to subsection 2.1, algae cells were prepared cell microscopic observation. The algae cells were diluted by adding 1 mL of the sample to 9 mL of distilled water, resulting in a final volume of 10 mL in triplicates. The samples were mixed using a vortex mixer to ensure accurate cell counting and measurements. 10 µL of the algae sample was introduced into the 0.1 µL JHS standard Thoma hemacytometer (0.100 mm depth × 1/400 mm^2^ width, Sunlead Glass Corp., Tokyo, Japan) for enumeration.

Another 10 µL of the algae sample was utilized to determine the algae cell size using an Olympus BX50 microscope with a ToupCam IP103100A camera and ToupView software for analysis. Both the counting and measurement were performed with a 40X lens magnification. A total of five samples were analyzed, with each sample subjected to ten replicates for cell number counts and twenty replicates for cell size measurements.

#### ii. Cultivation experiments to determine microalgae oxygen dynamics

The experiment was conducted under autotrophic conditions, as illustrated in Fig 1. 300 mL of CZ-M1 medium was prepared, based on the methodology previously described in subsection 2.1. The medium was autoclaved at 121 °C for 20 minutes using a Sanyo (Panasonic) MCV-B91S autoclave. During the algae cell washing procedure, 150 mL of *C. zofingiensis* and *H. lacustris* cells were centrifuged for 15 and 10 min at 10,000 rpm respectively due to their difference in size. The supernatants were removed after centrifugation and 0.05 % NaCl solution was added to the tubes. This centrifuged process and the addition of 0.05 % NaCl in the tubes were repeated three times. Following the washing procedure, 15 mL of 0.05 % NaCl solution was added to the tubes in preparation for the experiment.

**Fig 1:**
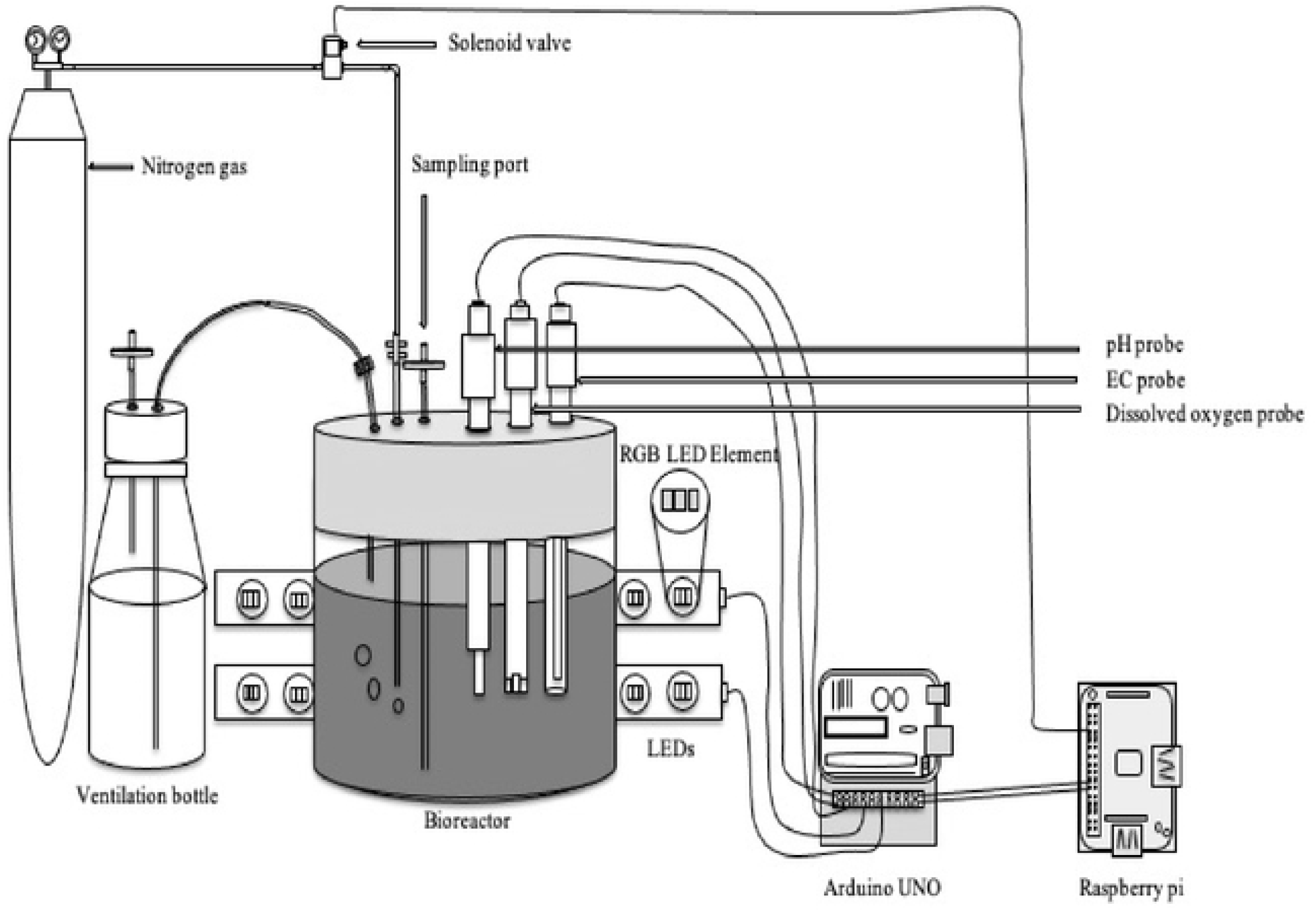
Schematic overview of the stirred tank reactor (STR). The cultivation of *C. zofingiensis* and *H. lacustris* was conducted under autotrophic growth conditions in glass flasks. A solenoid valve regulated the supply of nitrogen gas (N_2_) in the reactor. Light in the blue and red spectra was provided at different intensities by LEDs. Probes for measuring EC, DO, and pH were used and connected to an Arduino UNO microcontroller, with data recording managed via a Raspberry Pi.

A total of 15 mL of washed *C. zofingiensis* and *H. lacustris* cells was introduced into the reactor, which had the following dimensions: height 9.3 cm x diameter 8.9 cm, filled with autoclaved CZ-M1 medium. The medium was agitated with a magnetic stirrer set at 250 rpm to initiate the experimental procedure. The stirred tank reactor (STR) was placed in an incubator to maintain a constant temperature at 25 °C without aeration.

pH sensor (Horiba 9680S-10D) and electrical conductivity (EC) probe (Horiba 3552-10D) was connected to an Arduino Uno microcontroller via Atlas Scientific Ezo circuits. A dissolved oxygen (DO) probe (DFRobot, Gravity: SKU SE No. 237) was also utilized and connected to the controller. These probes were inserted into the reactor through holes in the silicon plug. The sensors were calibrated before starting the experiment.

A Python and Arduino programs were developed for the purpose of monitoring and controlling the system. Light was provided by using light-emitting diodes (LED) as an energy source. The LED can illuminate red, green and blue light in a single chip at different strengths as programmed. The LED chips were aligned and connected to the microcontroller.

The system was observed in both blue- and red-light environments, with light intensities of 20, 40, 60, and 80 µmol m^−2^ s^−1^, with 40 minutes of light exposure followed by 40 minutes of darkness per cycle for 24 hours. At the start of each cycle, the algae cells were exposed to nitrogen gas (N_2_) for 5 minutes without light to reduce the dissolved oxygen level. N_2_ was supplied from a cylinder, using a solenoid valve (MLV-3-1/8, 200 kPa, Takasago, Japan) connected to a relay module, which was controlled by the microcontroller. The study cycle was repeated at different light wavelengths and intensities for 2 days.

The Arduino microcontroller was connected to a Raspberry Pi 3 via USB cable for real-time data recording and analysis. The observed data of pH, EC and DO were sent from the microcontroller to the Raspberry Pi via serial communication at every 10 seconds to be recorded on a Secure Digital card.

6 mL of samples were collected daily from the reactor sampling port for algal absorbance analysis under each light condition. This experimental study was repeated three times for each light wavelength and for each algal culture.

### 2.3 Determination of biomass concentrations

Samples for algae biomass concentrations analysis were obtained from the cultivation experiments to determine microalgae oxygen dynamics experiment. During the study, it was not possible to obtain the requisite volume of liquid for the measurement of suspended solids (SS) at each sampling. Consequently, absorbance was measured using UV-VIS mini-1240, Shimadzu, Japan. While the SS was estimated from the result using the following equation developed from preliminary studies in our laboratory (*R* ² = 0.999):

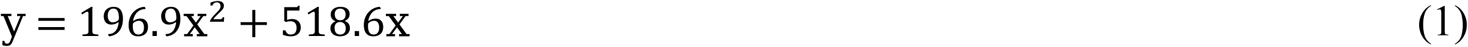

where *y* is estimated SS (mg L^−1^), *x* is the absorbance at 660 nm.

#### a) Oxygen production rate (*OPR*)

The *OPR* (µmol O_2_ gSS^−1^ min^−1^) was calculate d from the slope of the dissolved oxygen concentration over the last 40 minutes of the light phase 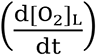, divided by the biomass concentration (C_b_) (eq.1).

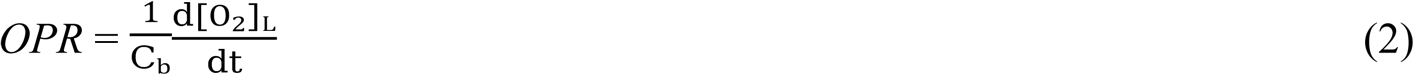

#### b) Oxygen consumption rate (*OCR*)

The *OCR* (µmol O_2_ gSS^−1^ min^−1^) was calculated from the slope of the dissolved oxygen concentration over the last 45 minutes of the dark phase 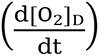, divided by the biomass concentration (*C*_b_) (eq.1).

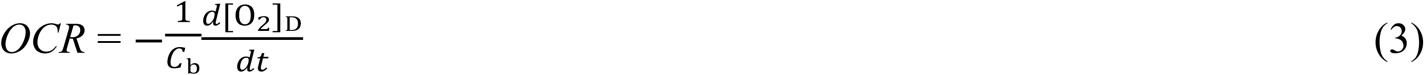

#### c) Microalgae net photosynthetic rate (*MNPR*)

The *MNPR* was calculated as the difference between *OPR* and *OCR* (µmol O_2_ gSS^−1^ min^−1^).

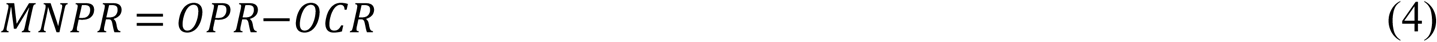

#### d) Photosynthetic quotient (*PQ*)

The *PQ* was determined as the ratio between the oxygen production rate (eq.2) and oxygen consumption rate (eq.3).

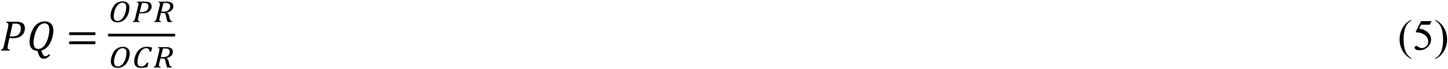

## 3. Results and discussions

### 3.1 Algae cell characteristics

Table 1 presents cell characteristic comparison of *C. zofingiensis* and *H. lacustris.* The surface area and volume were calculated based on the diameter of each cell assuming spherical shape of the algae. The results revealed that the average cell diameter of *H. lacustris* was larger than that of *C. zofingiensis*. During the cell size analysis, the maximum and minimum cell diameters for *C. zofingiensis* were 8.32 µm and 1.99 µm, respectively, while *H. lacustris* had a maximum and minimum cell diameter of 26.86 µm and 11.42 µm, respectively as shown in Fig 2. Additionally, average surface area and volume per cell of *H. lacustris* were about 9 and 24.1 times as large as *C. zofingiensis* cells. The algae cell sizes were with the ranges stated by other researchers [27,29].

**Fig 2:**
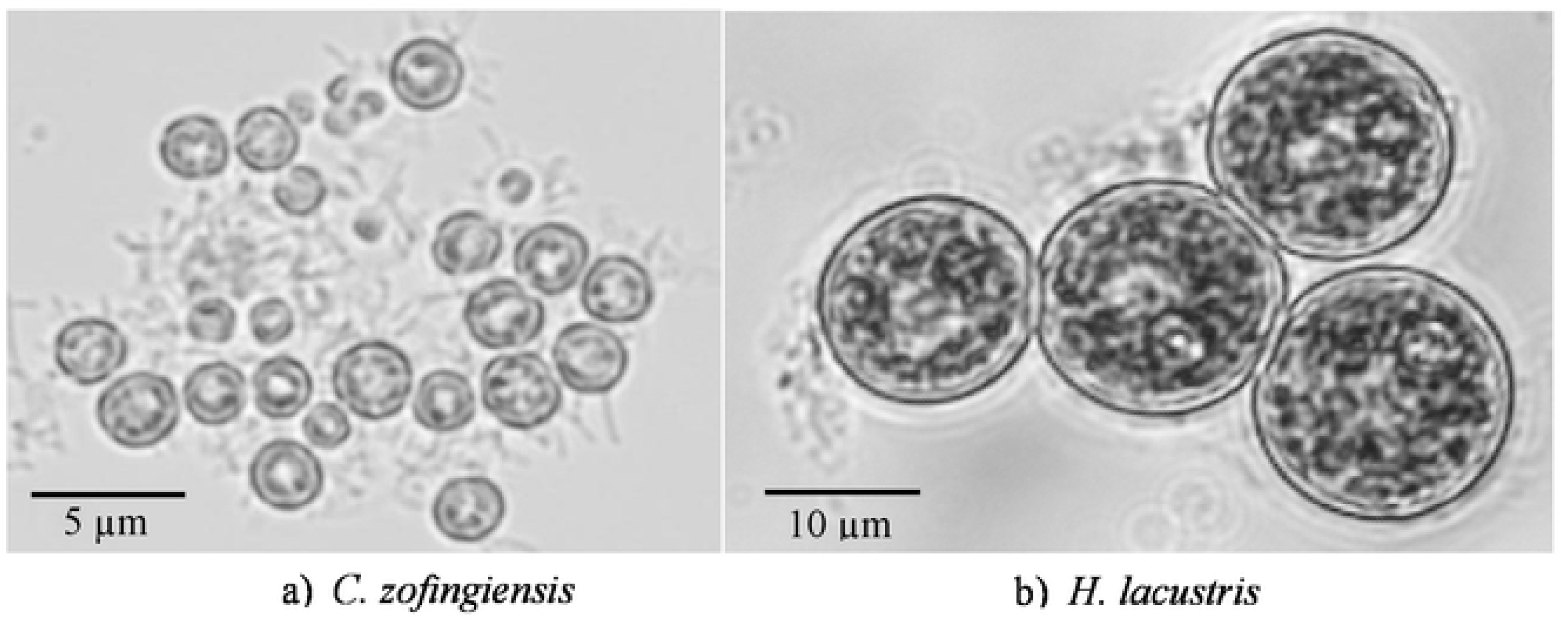
Microscopic images of *C. zofingiensis* (a) and *H. lacustris* (b). *C. zofingiensis* cells measured 1.99–8.32 µm in diameter, while *H. lacustris* cells measured 11.42–26.86 µm. Scale bars: 5 µm (a), 10 µm (b). The cells were viewed at 40X magnification with Olympus BX50 microscope with a ToupCam IP103100A camera and ToupView software for analysis. Thoma hemacytometer (0.100 mm depth × 1/400 mm2 width, Sunlead Glass Corp., Tokyo, Japan) was used to determine the algae cell size. Both algae were viewed at their exponential growth stages.

**Table 1:** Algae cell characteristics. Algae samples were obtained from the algae growth chamber. 1 mL of algae samples were added to 9 mL of distilled water, to make up 10 ml of working volume for each tube. The samples were vibrated for uniform mixture for algae cell counting and measurement. 10 µl of algae sample was injected into the 0.1 µl JHS standard Thoma hemacytometer and also 10 µl of algae sample was injected into the objective micrometer glass to measure the algae cell size. Both the algae counting and measurement was performed with 40X magnification. (a) cell size, surface area and volume (*n =* 150) were measured to determine the algae characteristic differences. (b) Data represents means ± SD (*n* = 5).

The smaller cell size of *C. zofingiensis* resulted in a larger surface-to-volume ratio, i.e. specific surface area, of 9.6 ± 6.4×10^−5^ m^2^ m^−3^ compared to 3.4 ± 2.1×10^−6^ m^2^ m^−3^ of *H. lacustris.* Assuming equal flux density at the plasma membrane, a larger specific surface area would facilitate more efficient transport of molecules and ions for metabolism. This enhanced surface area could improve the uptake of essential nutrients and the discharge of waste products. In contrast, the smaller specific surface area of *H. lacustris* may limit membrane transport, thereby contributing to its lower specific growth rate. Furthermore, it could also contribute to the less protection of the intracellular environment from changes in the external environment.

### 3.2 Effect of different light wavelengths and intensities on *c. zofingiensis* and *h. lacustris* oxygen production and consumption rates per biomass

#### 3.2.1 Algae Oxygen Production (OPR) per Biomass

The oxygen production rates of *C. zofingiensis* and *H. lacustris* were evaluated in different light wavelengths and intensities with the results shown in Fig 3 (a) and (b) respectively. During the cultivation of *C. zofingiensis*, the OPR decreased under blue-light conditions, from 10 to 1.4 µmol O_2_ gSS^−1^ min^−1^ as the light intensities were changed from 20 to 80 µmol m^−2^ s^−1^. Similarly, under red-light condition, the OPR were declined from 1.83 to 0.67 µmol O_2_ gSS^−1^ min^−1^ over the same range of light intensity. These changes observed in Fig 3 (a) and (b) indicate the effect of low light intensity in OPR. The comparison between different light wavelengths also shows that, under red-light conditions at 20 µmol m^−2^ s^−1^ the OPR was 1/5 compared to the OPR exhibited by *C. zofingiensis* under blue-light conditions at the same intensities. Similarly at 80 µmol m^−2^ s^−1^ under red-light condition, there was further decrease in OPR, which was 2.1 times lower compared to blue-light conditions at same intensity.

**Fig 3:**
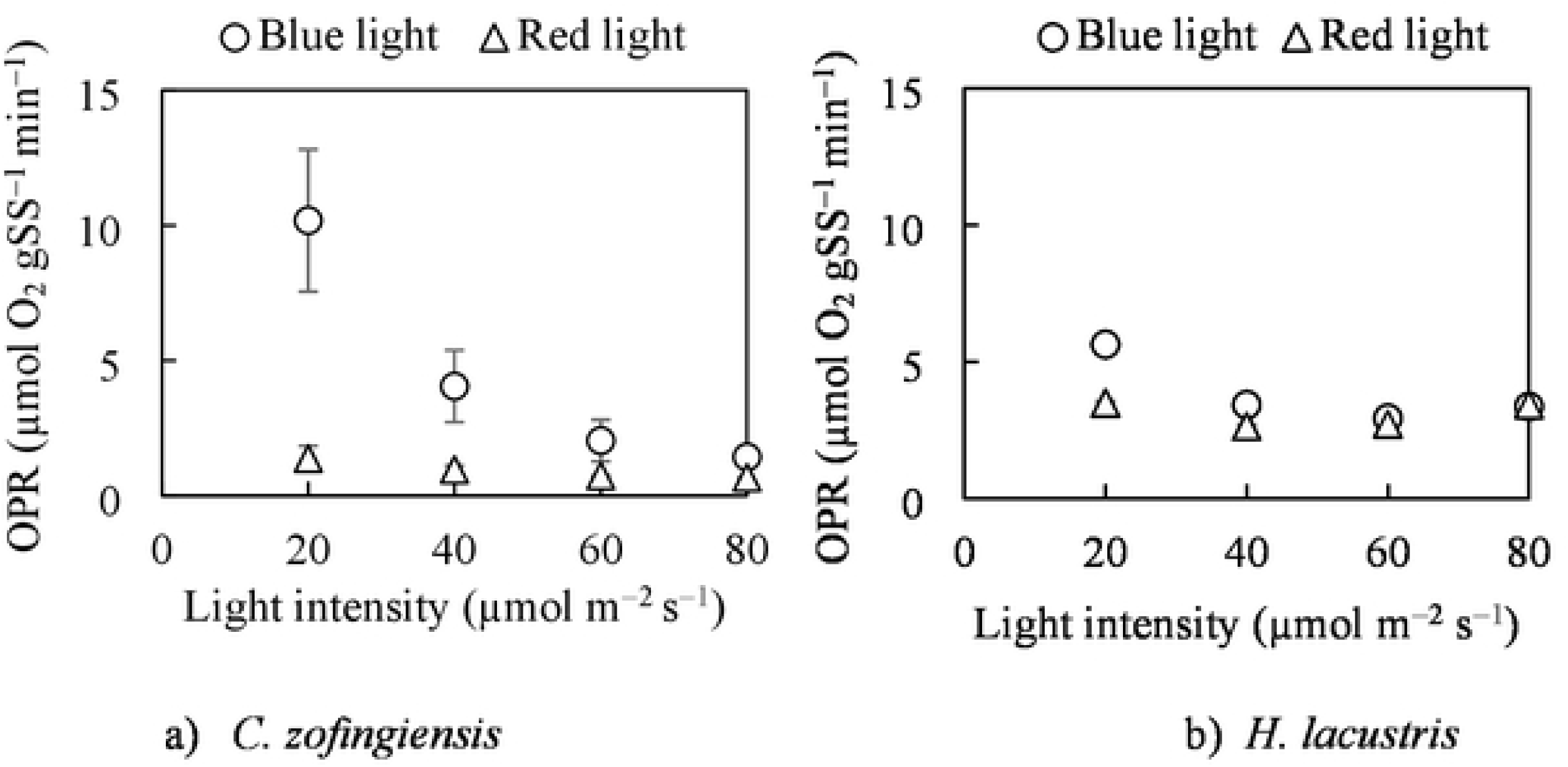
Oxygen production rate (OPR) per biomass under different light wavelengths and intensities. This shows *C. zofingiensis* (a) and *H. lacustris* (b) Oxygen production under blue and red light 20–80 µmol m^−2^ s^−1^. Algae cell oxygen consumption was measured for a 10-day period using a dissolved oxygen probe. Comparing oxygen production rates at different light wavelengths and intensities. Error bar represents standard deviation (*n* = 3).

*H. lacustris* cells exhibited a decrease in OPR with change in light intensity in both blue and red-light conditions in Fig 3 (b). Under blue-light conditions at 20 µmol m^−2^ s^−1^, the highest OPR was 5.63 µmol O_2_ gSS^−1^ min^−1^ and this value decreased with change in light intensity to 3.41 µmol O_2_ gSS^−1^ min^−1^ at 80 µmol m^−2^ s^−1^. Furthermore, under red-light conditions, the OPR were withing range of 3.49–2.64 µmol O_2_ gSS^−1^ min^−1^ with the highest value observed at 20 µmol m^−2^ s^−1^. These OPR values were observed to decrease with change in light intensities.

Comparison between the OPR of these algae species revealed that *H. lacustris* exhibited OPR values which was 1.6 times smaller compared to *C. zofingiensis* at 20 µmol m^−2^ s^−1^. The OPR exhibited by *H. lacustris* can be attributed to the increased cell size and cell surface which affected its oxygen molecules transport (Table 1). Furthermore, under the red-light conditions, *H. lacustris* also exhibited OPR range which were higher compared to *C. zofingiensis* under the same conditions due to the adaptability of *H. lacustris* under red-light.

Under identical photon flux densities, as shown in Table S1, the total energy input from blue light (470 nm) was approximately 40 % higher than from red light (660 nm). For example, at 80 µmol m^−2^ s^−1^, the radiant power was 20.4 W·m^−2^ under blue illumination compared with 14.5 W·m^−2^ (Table S2) under red light. This difference arises from the higher photon energy (4.23 × 10⁻¹⁹J photon^−1^) of blue light relative to red (3.01 × 10^−19^ J photon^−1^) (Table S2). The additional optical energy enhances PSII excitation and electron flow [30]. resulting in the increased oxygen production rate and carotenoid biosynthesis observed in *C. zofingiensis* and *H. lacustris* in Fig 3 (a) and (b).

#### 3.2.2 Oxygen consumption rates (OCR) per biomass

In the absence of photosynthesis, algae engage in cellular respiration to meet their energy needs. During this process, cellular organic compounds, such as glucose, are broken down in the presence of oxygen to release energy. This process consumes oxygen and produces carbon dioxide and water as by-products [28].

During the OCR analysis, an increase in the dissolved oxygen (DO) was observed in the reactor during the dark phase (zero light intensity) in both algae cultures. It was postulated that this increase in DO concentration had been due to oxygen excretion from the internal cell space of algae or re-circulation of displaced oxygen during the N_2_ gas supply (data not shown). This resulted in an equilibrium state between the reactor headspace and the medium. The DO increase in the dark phase was in positive values and based on eq. 3, the negative OCR values recorded signified oxygen increase consumption.

In Fig 4 (a) *C. zofingiensis* showed a negatively large OCR at 20 µmol m^−2^ s^−1^ as –7.08 µmol O_2_ gSS^−1^ min^−1^ in blue-light conditions due to the increase supply of oxygen. The oxygen utilization in this environment resulted in specific growth rate of 0.52 d^−1^ as shown in Fig 5. The OCR was observed to vary in response to changes in blue-light intensity. At 80 µmol m^−2^ s^−1^ the OCR was approximately –0.70 µmol O_2_ gSS^−1^ min^−1^ which was less than the observations at 20 µmol m^−2^ s^−1^ of light intensity, due to reduced growth rate and lesser negative OCR. The result showed that change in light intensity had an effect on algae growth rate and oxygen consumption. Under red-light conditions at 20 µmol m^−2^ s^−1^, the OCR during the dark phase increased to –0.63 µmol O_2_ gSS^−1^ min^−1^. At this intensity, as shown in Fig 5, the specific growth rate reached 0.14 d^−1^, which was the highest observed values under the red-light condition with *C. zofingiensis*. However, this growth rate was still lower compared to that observed under blue-light conditions at the same light intensity. Further change in light intensity under red-light conditions, showed more less negative OCR (–0.12 µmol O_2_ gSS^−1^ min^−1^), while *C. zofingiensis* exhibited lower growth rate to 0.01 d^−1^ at 80 µmol m^−2^ s^−1^.

**Fig 4:**
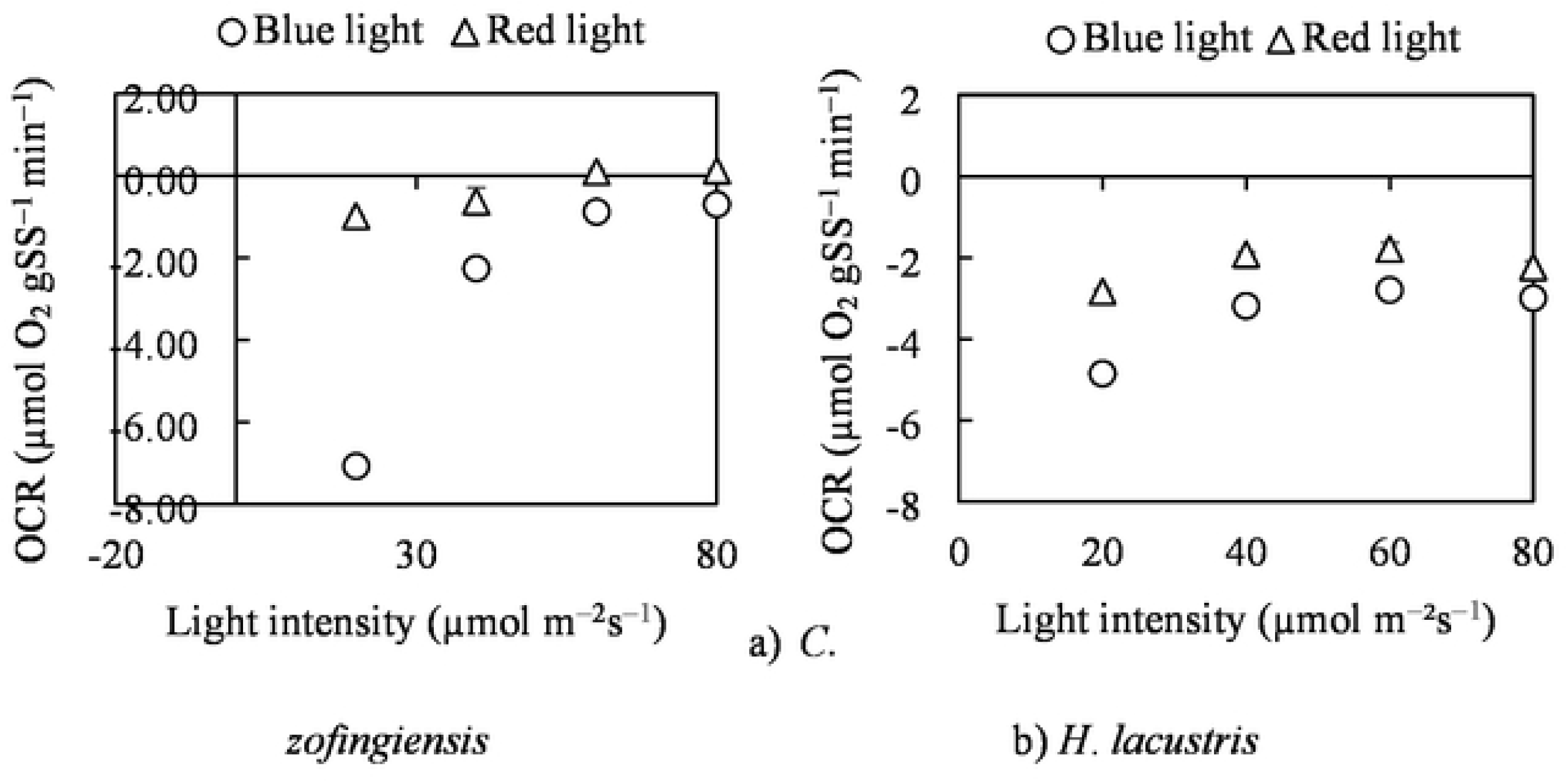
Oxygen consumption rate (OCR) per biomass under different light wavelengths and intensities. This shows *C. zofingiensis* (a) and *H. lacustris* (b) Oxygen consumption under blue and red light 20 – 80 µmol m^−2^ s^−1^. Algae cell oxygen consumption was measured for a 10-day period using a dissolved oxygen probe. Comparing oxygen consumption rates at different light wavelengths and intensities. Error bar represents standard deviation (*n* = 3)

**Fig 5:**
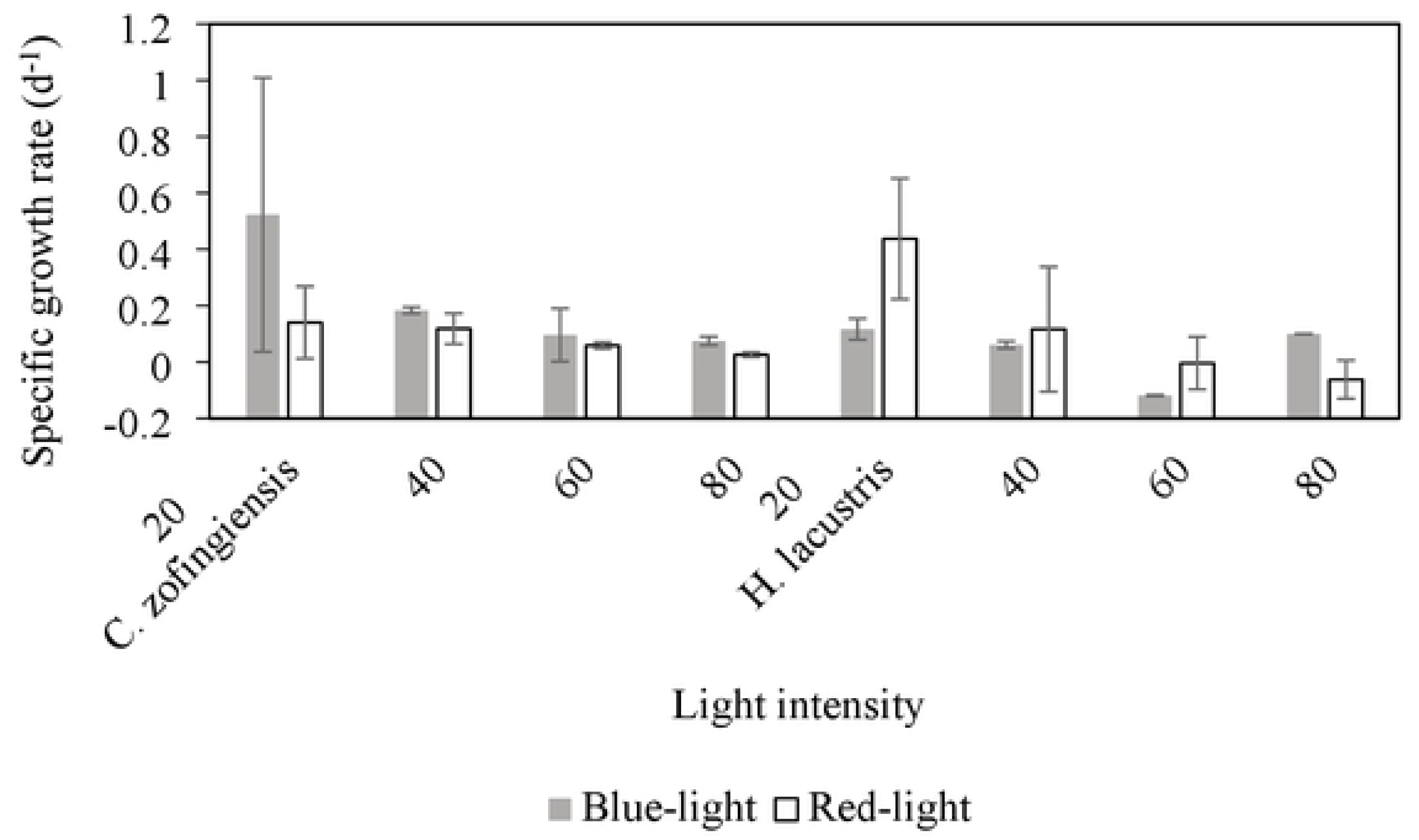
Algae specific growth rate under different light wavelengths and intensities. The effect of different light wavelengths (blue and red) at 20–80 µmol m^−2^ s^−1^ on *C. zofingiensis* (left) and *H. lacustris* (right) specific growth rate. The algae growth was measured for a 10-day period using a photospectrometer. Error bar represents standard deviation (*n* = 3).

In Fig 4 (b) *H. lacustris* responded to change in different light wavelengths and intensities. Under blue-light conditions at 20 µmol m^−2^ s^−1^, *H. lacustris* exhibited OCR of – 6.42 µmol O_2_ gSS^−1^ min ^−1^ and exhibited a growth rate of 0.12 d^−1^. Further change in light intensity, from 20–80 µmol m^−2^ s^−1^, led to less large OCR and decreased growth rate, as shown in Fig 4 (b) and Fig 5. Additionally, under red-light conditions, lower OCR values were observed compared to those under blue-light conditions. The highest OCR under red-light was –2.84 µmol O_2_ gSS^−1^ min^−1^, recorded at 20 µmol m^−2^ s^−1^ which also corresponded to an increased specific growth rate of 0.43 d^−1^ compared to the blue-light conditions. We speculated that blue-light wavelength could have put *H. lacustris* in a stress condition, which affected its growth and delayed cell division [31]. A study that compared the growth of *H. lacustris* under different light wavelengths and intensities also found that, its performance under red-light at 18 µmol m^−2^ s^−1^ was greater compared under blue and white-light at the same light intensity [32].

*C. zofingiensis* exhibited a significant increase in both OCR and specific growth rate across the light intensity range of 20–80 µmol m^−2^ s^−1^ under the blue-light conditions compared to its performance under red-light and the responses of *H. lacustris* in similar conditions. *C. zofingiensis* cell size also had influenced its increased growth rates compared to *H. lacustris* due to the later increased cell surface area and smaller cell size for streamlined nutrient transportation and absorption. The increase in growth rate of *H. lacustris* at 20 µmol m^−2^ s^−1^ under red-light condition indicates its physiological adaptation. The high oxygen consumption observed in *C. zofingiensis* under blue-light conditions at low light intensity suggests elevated hydroxyl radical production, potentially supporting carotenogenesis and astaxanthin biosynthesis as previously reported [13]. This indicates that *C. zofingiensis* is an economically viable microalgae for the cost-effective strategy in the production of carotenoids at low light intensity.

### 3.3 Effect of different light wavelengths and intensities on microalgae net photosynthetic rate (MNPR) and photosynthetic quotient (PQ)

Table 2 shows the results of investigations conducted to determine the MNPR and the PQ, as part of microalgae cell characteristics as shown in eq. 4 and 5, respectively. The MNPR is the difference between the oxygen production rate and the oxygen consumption rate [10] while the PQ as mentioned earlier deals with the ratio of oxygen production and consumption rates [9]. these are important parameters in determining the reduction of CO_2_ to carbohydrate when all the electrons released from water splitting in PSII are utilized. However, when electrons are drawn from photosynthetic chemical reactions to other reductive sinks, more oxygen will be produced than oxygen consumed, and PQ will be greater than 1.0 [33].

**Table 2:** Effect of different light wavelengths and intensities on MNPR and PQ of *C. zofingiensis* and *H. lacustris* in both blue- and red-light environment. MNPR and OPR comparison of *C. zofingiensis* and *H. lacustris* were carried in both blue and red-light environment in 20–80 µmol m^−2^ s^−1^ of light intensity for a 10-day period. The MNPR was calculated from the difference between the OPR and OCR while PQ was calculated from the ratio between OPR and OCR.

In the presence of blue-light, the net photosynthetic rate of *C. zofingiensis* was found to be highest at 20 µmol m^−2^ s^−1^ with a rate of 3.10 ± 12.41 µmol O_2_ gSS^−1^ min^−1^ showing a large gap between oxygen production and consumption rate. This rate was decreased with change in light intensity to approximately 4.2 times at 80 µmol m^−2^ s^−1^. Furthermore, in the red-light conditions, the MNPR observed were lower compared to the blue-light conditions at each light intensity. In these conditions (red-light) the MNPR was observed to increase with change in light intensity at 20–40 µmol m^−2^ s^−1^ indicating an increased absolute difference between oxygen production and consumption. At 60–80 µmol m^−2^ s^−1^, the oxygen production rate was observed to decline creating an oxygen deficient condition. This phenomenon underscores the positive effect of blue-light at lower light intensity. *H. lacustris* exhibited a poor MNPR compared to *C. zofingiensis* under blue-light conditions with the highest MNPR (2.38 µmol O_2_ gSS^−1^ min^−1^) observed at 20 µmol m^−2^ s^−1^. In the red-light conditions, The MNPR were observed to increase with change in light intensity and were higher compared to the rates in the blue-light conditions. Generally, the MNPR observed with *C. zofingiensis* under blue and red-light conditions were higher compared to MNPR with *H. lacustris* in both conditions.

Furthermore, the observed decline in MNPR with change in light intensity under blue-light conditions suggests increased utilization of blue-light energy for biomass accumulation by *C. zofingiensis* at intensities below 80 µmol m^−2^ s^−1^. In contrast, the increase in MNPR under red-light (*C. zofingiensis*), blue and red-light (*H. lacustris*) shows poor utilization of produced O_2_ for biomass accumulations as shown in Fig 5.

Under blue-light conditions, *C. zofingiensis* exhibited PQ values greater than 1 as light intensity increased from 20 to 80 µmol m^−2^ s^−1^, indicating enhanced photosynthetic efficiency under this light wavelength. The highest PQ value of 2.3 was recorded at 60 µmol m^−2^ s^−1^. In contrast, under red-light conditions, elevated PQ values were observed only at lower intensities (20–40 µmol m^−2^ s^−1^), suggesting higher oxygen production within that range. Under blue-light conditions, *H. lacustris* exhibited PQ values ranging from 0.9 to 1.1 as light intensity increased from 40 to 80 µmol m^−2^ s^−1^. These values were lower than those observed under red-light conditions at the same intensities. Under red-light conditions, PQ values were above 1.0 across the 20–80 µmol m^−2^ s^−1^ range, with the highest PQ of 1.5 recorded at 60 and 80 µmol m^−2^ s^−1^, indicating enhanced oxygen production under red light at high light intensity.

The comparison of oxygen production and consumption rates, as well as responses to different light wavelengths and intensities, between *C. zofingiensis* and *H. lacustris* revealed distinct differences in their photosynthetic capacities. *C. zofingiensis* exhibited higher PQ values under blue-light conditions compared to *H. lacustris* under both blue and red light, as well as compared to the PQ value reported for *Chlorella sorokiniana* at 1500 µmol m^−2^ s^−1^[10]. As previously observed, there is a strong correlation between photosynthetic efficiency and the synthesis of lipids and carotenoids [20]. This suggests that *C. zofingiensis* possesses enhanced photosynthetic performance, particularly in oxygen evolution relative to oxygen uptake, which could contribute to the accumulation of high-value biomolecules. Furthermore, the observed high MNPR and PQ values across different blue-light intensities highlight the potential of *C. zofingiensis* as a promising source for efficient oxygen production.

### 3.4 Effect of different light wavelengths and intensities on the culture media pH of *C. zofingiensis* and *H. lacustris*

The pH profiles of *C. zofingiensis* and *H. lacustris* as shown in Fig 6 were monitored under various light wavelengths and intensities, in CZ-M1 medium with initial pH range of 6.8–7.0 as described in subsection 2.1. Under both blue and red-light conditions, the pH values increased in response to changes in light intensity during the algae culture in Fig 6 (a) and (b). Under blue-light conditions with *C. zofingiensis*, the effect of change in light intensity, resulted in a subsequent rise in pH values from 7.0 at 20 µmol m^−2^ s^−1^ to 8.1 at 80 µmol m^−2^ s^−1^ in Fig 6 (a). Conversely, under red-light conditions, increase in pH were also observed with changes in light intensity, with the highest as 9.5–9.6 at 80 µmol m^−2^ s^−1^ in Fig 6 (b). During the *H. lacustris* culture, pH increase was also observed with change in light intensity as shown in Fig 6 (c) and (d). These pH values were higher compared to the observed pH with *C. zofingiensis* medium utilizing the same light wavelengths and intensities. *H. lacustris* media pH increased from 7.0 initial pH to 9.5 and 11.8 at 80 µmol m^−2^ s^−1^, under the blue and red-light conditions respectively.

**Fig 6:**
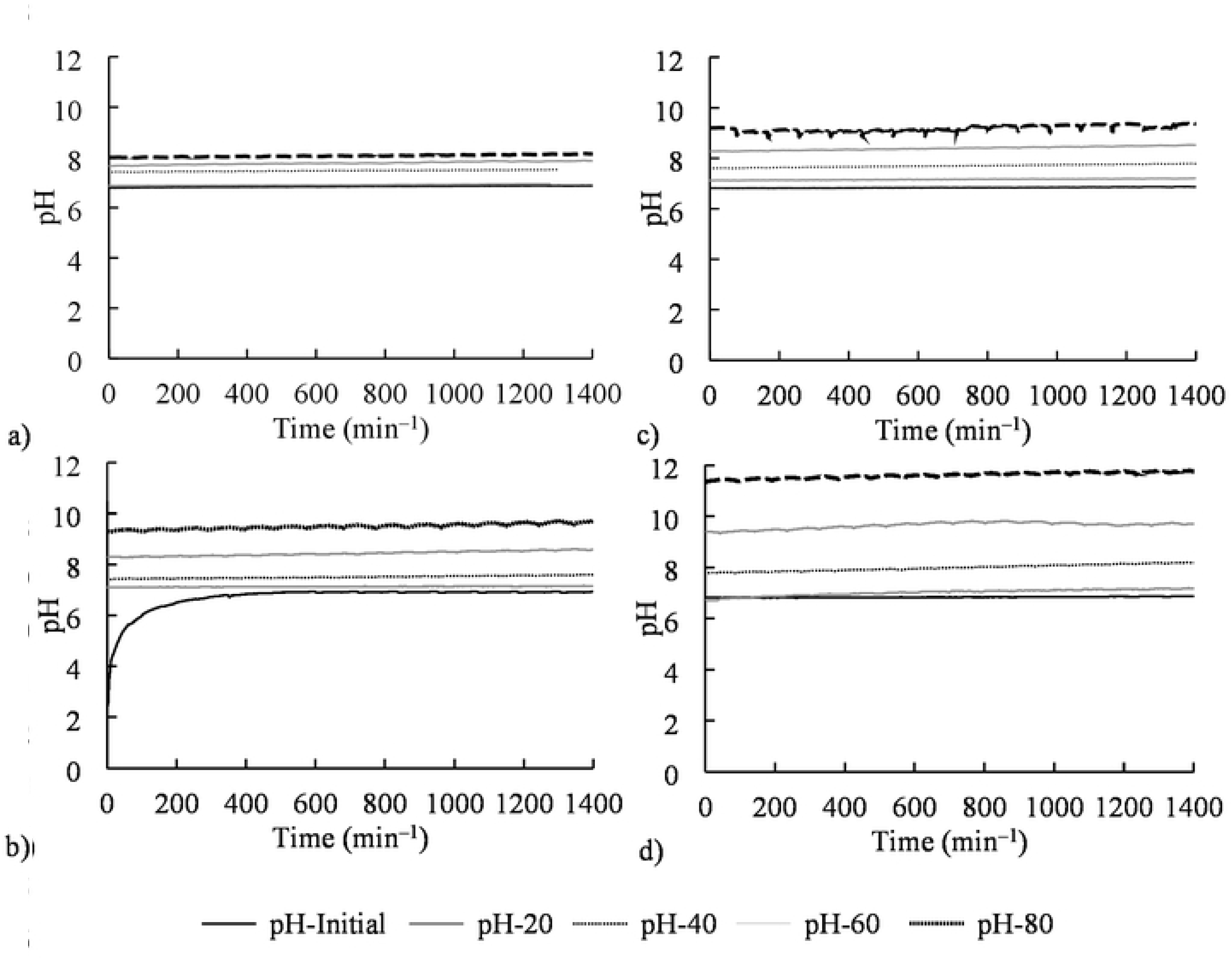
The effect of different wavelengths and light intensities on the culture media pH of *C. zofingiensis* and *H. lacustris*. The effect of light wavelengths and intensities (20–80 µmol m^−2^ s^−1^) on algae culture environment were measured using a pH probe. Comparison of change in pH under different light wavelengths and intensities. *C. zofingiensis* a) blue- and b) red-light, *H. lacustris* c) blue- and (d) red-light. Error bar represents standard deviation (n = 3).

The results indicate that pH variations were influenced by different light wavelengths and intensities during autotrophic condition of the algae. Generally, under blue-light conditions, lower pH values were observed compared to red-light conditions in both algae species (Fig 6). This phenomenon is attributed to the red-light long wavelength effect on algae cells. The reduction of CO_2_ resulted in a shift in chemical equilibrium, leading to a decrease in the concentration of H^+^ ions and an increase in pH, thereby rendering the medium more alkaline. Furthermore, the lower pH values observed with *C. zofingiensis* media compared to *H. lacustris* media may be attributed to the increased production and utilization of produced oxygen in both blue and red-light conditions for growth. This suggests that the balance between oxygen production and consumption in the medium were relatively stable and renders the media pH suitable for algae growth [34]. The increased pH (9.5–11.8) in *H. lacustris* media had affected its growth. This observation is relative to the suggestion that increase in pH could lead to poor fixation and utilization of absorbed CO_2_ [2].

## 4 Conclusion

This study characterized the cell size and oxygen exchange behavior of *C. zofingiensis* and *H. lacustris* under different light wavelengths and intensities in autotrophic conditions. The real-time monitoring approach effectively measured oxygen production and consumption rates without disrupting algae growth. Both species responded differently to variations in light wavelength and intensity. Blue-light supported higher oxygen production rates in both algal species compared to red-light. *C. zofingiensis*, due to its smaller cell size and higher surface area-to-volume ratio, exhibited significantly greater oxygen production (OPR) and consumption (OCR) rates particularly at light intensities below 60 µmol m^−2^ s^−1^. The blue-light conditions also supported *C. zofingiensis* high growth rates and stable pH levels. In contrast, *H. lacustris*, with its larger cell size, showed lower OPR and OCR efficiency under the same conditions but achieved high growth under red-light only at 20 µmol m^−2^ s^−1^. The results highlight the role of cell size and blue-light at low intensity in promoting oxygen production and growth rate in *C. zofingiensis*, positioning it as a cost-effective candidate for combined oxygen generation.

## Acknowledgement

The authors are thankful to the Cabinet Office grant in aid "the Advanced Next-Generation Greenhouse Horticulture by IoP (Internet of Plants)" and "Evolution to Society 5.0 Agriculture Driven by IoP (Internet of Plants)" Japan. Additionally, we also wish to thank, the Ministry of Education, Culture, Sports, and Technology (MEXT) in collaboration with Okayama University, Japan.

## Author contribution

Ugwu Chigozie: Performed the experiments, analyzed and interpreted the data, and drafted the paper. Hideaki Nagare: Conceived and designed the experiments, analyzed and interpreted the data, contributed reagents, and supervised the experiments. Ayumi Hashiguchi: Reviewed and edited the paper.

## Availability of data and materials

All datasets on which the conclusions of the manuscript rely are presented in the main paper.

## Conflict of interest statement

The authors affirm that they have no known financial or interpersonal interest that would have appeared to have an impact on this study.

## Funding statement

This work was supported by Cabinet Office grant in aid, “the Advanced Next-Generation Greenhouse Horticulture by IoP (Internet of Plants)” and “Evolution to Society 5.0 Agriculture Driven by IoP (Internet of Plants),” Japan.

## Notes

### Competing Interest Statement

The authors have declared no competing interest.

